# An expedited qualitative profiling of free amino acids in plant tissues using liquid chromatography-mass spectrometry (LC-MS) in conjunction with MS-DIAL

**DOI:** 10.1101/2024.06.28.601148

**Authors:** Anish Kaachra, Anish Tamang, Vipin Hallan

## Abstract

The estimation of relative levels of amino acids is crucial for understanding various biological processes in plants, including photosynthesis, stress tolerance, and the uptake and translocation of nutrients. A wide range of liquid chromatography (LC; HPLC/UHPLC) -based methods is available for measuring the quantity of amino acids in plants. Additionally, the coupling of LC with mass spectrometry (MS) significantly enhanced the robustness of existing chromatographic methods used for amino acid quantification. However, accurate annotation and integration of mass peaks can be challenging for plant biologists with limited experience in analyzing MS data, especially in studies involving large datasets with multiple treatments and/or replicates. Further, there are instances when the experiment demands an overall view of the amino acids profile rather than focusing on absolute quantification. The present protocol provides a detailed LC-MS method for obtaining a qualitative amino acids profile using MS-DIAL, a versatile and user-friendly program for processing MS data. Free amino acids were extracted from the leaves of control and Tomato leaf curl Palampur virus (ToLCPalV)-infected *Nicotiana benthamiana* plants. Extracted amino acids were derivatized and separated using UHPLC-QTOF, with each amino acid subsequently identified by aligning mass data with a custom text library created in MS-DIAL. Further, MS-DIAL was employed for internal standard-based normalization to obtain a qualitative profile of 15 amino acids in control and virus-infected plants. The outlined method aims to simplify the processing of MS data to quickly assess any modulation in amino acid levels in plants with a higher degree of confidence.

## Introduction

Amino acids play a crucial role in the growth and development of plants. Analyzing the amino acid profile provides valuable insights into the rate of various processes and the metabolic status of plants (Trovato et al., 2021; Foyer et al., 2003). Amino acids exist in plants both in free form and bound to other metabolites. The pool of free amino acids plays a crucial role in various physiological processes including growth regulation, nitrogen uptake and translocation, coordination of carbon and nitrogen metabolism, seed development, and nutritional value. Numerous studies indicate a positive correlation between the accumulation of specific amino acids and certain biological processes. For example, proline (Pro) accumulates under stress conditions (Hayat et al., 2012), while the absolute amounts of glutamine (Gln), glycine (Gly), and serine (Ser) are influenced by nitrogen assimilation and photorespiration (Foyer et al., 2003). The Gly/Ser ratio indicates the carbon flux through photorespiration (Novitskaya et al., 2002), and the levels of lysine (Lys) and methionine (Met) reflect the nutritional quality of seeds (Yang et al., 2020; Whitcomb et al., 2020).

Liquid chromatography (HPLC/UHPLC) with pre-column derivatization is a widely used technique for amino acid analysis due to its high sensitivity and resolution. Notably, an idle derivatization agent not only improves chromatographic separation but also facilitates specific detection methods, such as UV absorption or fluorescence-based detection of amino acids (Ferre et al., 2019). Commonly used derivatization reagents include ortho-phthalaldehyde, 9-fluorenylmethyl chloroformate (FMOC), dansyl chloride, ninhydrin, and aminoquinolyl-N-hydroxysuccinimidyl carbamate (AQC) (Dahl-Lessen et al., 2018). While liquid chromatography (LC) appears to be a straightforward method for amino acid quantification, the assay often presents challenges that can be time-consuming and intricate to resolve. For instance, prolonged run/analysis times, incomplete derivatization, and the appearance of extra peaks add complexity to the analyses, particularly when dealing with crude plant extracts (Dahl-Lassen et al., 2018; Eid et al., 2022).

Technological advancements have led to the integration of LC with various MS techniques such as ESI-TOF MS, single quadrupole MS, triple quadrupole MS, or orbitrap MS to evolve faster and more reliable method for amino acid analysis (Dziągwa-Becker et al., 2015; Nemkov et al., 2015). In addition to enhanced sensitivity, LC-MS based methods make it easier to eliminate false annotation of LC peaks (Thiele et al., 2012; Guo et al., 2019; Eid et al., 2022). While derivatization agents may enhance the ionization of amino acids, MS methods have also been developed to allow the direct analysis of amino acids without derivatization (Thiele et al., 2019; Chen et al., 2021). Despite its many advantages, MS does have certain limitations including sensitivity to contaminants, variability in the ionization of amino acids, and the challenge of limited quantification without authentic standards or the necessity of expensive isotopically labeled standards. The complexity of MS instruments and the data generated can pose challenges for users without specialized training. Additionally, the post-run identification of individual amino acids becomes perplexing, especially when they are bound with a derivatization agent. In many instances, plant researchers opt for obtaining a comprehensive overview of the amino acid profile rather than focusing on absolute quantification. This inclination is particularly evident in investigations that include multiple samples, treatments, and replicates.

A variety of tools are accessible online to analyze mass data, each designed to address specific research needs. Platforms such as Galaxy, MS-DIAL, and MetaboAnalyst, known for user-friendliness, along with specialized tools like XCMS Online and MZmine 2, enable researchers to effectively process, analyze, and interpret mass spectrometry data (Peralbo-Molina et al., 2022). Out of these, MS-DIAL (http://prime.psc.riken.jp/compms/msdial/main.html) stands out as a freely available universal program for processing MS data (Tsugawa et al., 2015). Since its introduction, MS-DIAL has gained significant attention accumulating over 2300 citations as of July 2024. In addition to its user-friendly graphic user interface, MS-DIAL offers plenty of features including spectral deconvolution for LC/MS and data-independent MS/MS, simplified criteria for peak identification, and support for all processing steps from raw data import to statistical analysis (Perez de Souza et al., 2021). It is important to highlight that precise metabolite annotation from mass spectra represents the most challenging aspect of MS data processing. Information on accurate mass (MS), fragmentation pattern (MS/MS), and retention time (RT) of reference standards significantly assist in the identification of metabolite from the mass data. However, the quantity of compounds within spectral libraries and those for which reference standards are accessible is limited. Also, variation in MS/MS or RT across various instrument platforms and laboratories may lead to spurious identification. Therefore, a customized library containing a combination of independent parameters such as MS, MS/MS, and RT of a reference standard measured under identical analytical conditions would significantly enhance the accuracy of metabolite identification in the sample.

In the present method, free amino acids were extracted from control and ToLCPalV-infected *N. benthamiana* at different time points post-infection. Extracted amino acids derivatized with AQC were separated using reverse phase UHPLC, and their respective mass spectra were acquired using QTOF-MS. Subsequently, a custom library containing mass information (combined mass of amino acid standards and AQC) and RT was prepared to facilitate post-run identification of amino acids using MS-DIAL. The annotated amino acids underwent additional processing steps, including normalization with glycine-1-13C as an internal standard, curation of individual mass chromatographs, and performing principal component analysis within MS-DIAL itself. These features substantially improved the accuracy of acquiring comparative amino acid profiles across plant samples understudy. Notably, the application of LC-MS in the quantification of amino acids with or without derivatization has been reported by several other authors. In one such study, Dahl-Lassen et al., (2018) described a method for high-throughput analysis of AQC-derivatised amino acids in plant samples using UHPLC-single quadrupole MS (Dahl-Lassen et al., 2018). However, these studies often lack detailed post-run methodology for interpreting MS data. Therefore, we aim to provide a simple and quick method to obtain qualitative amino acid profiles in plant samples circumventing major limitations associated with the interpretation of LC-MS data. The described method would aid plant biologists and researchers with basic knowledge of MS to obtain a snapshot of modulations in amino acid levels with a higher degree of confidence.

## Method

### Extraction of free amino acids from plant sample

*Nicotiana benthamiana* plants were grown in an insect-free growth chamber at 24°C with 70% relative humidity under a 16-hour light and 8-hour dark cycle. Infectious clones of Tomato leaf curl Palampur virus (ToLCPalV) DNA-A and DNA-B were introduced to the plants at the 2-3 leaf stage using *Agrobacterium tumefaciens*-mediated inoculation. The infiltration process involved applying a vacuum by pressing a fingertip on the ventral side of the leaf and introducing the *Agrobacterium* suspension using a 1 ml needleless syringe. The infiltrated plants were then maintained in the growth chamber. Plants treated only with the inoculation buffer served as controls. Leaf samples (100 mg) were collected from both control and ToLCPalV infected plants at different time points i.e. 24 h, 48 h, 72 h, 7 d, and 15 d (h, hours; d, days) post-infection. For extraction of free amino acids, leaf samples were crushed into fine powder in liquid nitrogen and suspended in 500 µl of 10 mM HCl. The clear supernatant obtained after centrifugation was collected as an amino acid extract.

### Derivatisation of amino acids

Amino acid derivatization was performed essentially as described by the manufacturer’s instructions (Waters, USA). Briefly, 10 µl of the extract was mixed with 20 µl of aminoquinolyl-N-hydroxysuccinimidyl carbamate (AQC) and 70 µl of borate buffer followed by incubation at 55° C for 10 min. The mixture was filtered through a 0.25 µm filter before analysis.

### Liquid Chromatography

Derivatized amino acids were separated on an Agilent Infinity II UHPLC (Agilent Technologies, USA) and detected with a diode array detector (DAD) with separation column, ACCQ-TAG™ ULTRA C18 (2.1 mm X 100 mm, 1.7µm). The water (A) and acetonitrile (B) each with 0.1% formic acid were used as a mobile phase in the maintained gradient system. A gradient elution method by varying the percentage of mobile phases A and B as 0 - 0.54 min, 99.90% A; 0.54-5.74 min, 90.0% A; 5.74-7.74 min, 78.80% A; 7.74-8.04 min, 40.40% A; 8.04-8.64 min, 40.40% A, 8.64-8.73 min, 99.90% A, 8.73-9.50 min, 99.90% A, 9.50-10.20 min, 99.90% A, and 10.20-11.00 min, 99.90% A, resulted in the separation of amino acids. Other UHPLC parameters were set as column temperature: 55 °C; injection volume: 2 µl; flow rate: 0.700 ml/min; and absorption wavelength of 260 nm

### Mass Spectrometry

The mass determination was performed using a high-resolution 6560 ion mobility quadrupole time-of-flight (IM-QTOF) LC-MS system (Agilent Technologies, USA) equipped with electrospray ionization (ESI) in positive mode. The drying gas flow rate was adjusted to 10.0 L/min, maintaining a constant gas temperature of 340 °C and a gas pressure of 35 psi. Additionally, the capillary voltage was set to 3500 V, accompanied by a fragmentor voltage of 390 V. The mass spectrometer scanned within the range of 100–1700 m/z at a rate of 1.50 spectra/s.

### Workflow for the qualitative profiling of amino acids using MS-DIAL

1. Download and install MS-DIAL software from the link (https://systemsomicslab.github.io/compms/msdial/main.html). All the data processing described in the present protocol is done with MS-DIAL version 4.9.221218. It is important to mention that MS-DIAL version 5 has been recently introduced with a couple of advanced features. Nevertheless, we used MS-DIAL 4 which effectively yielded the comparative amino acid profile as aimed in the present protocol.
2. Convert the vendor mass data files (.d files in case of Agilent) to .abf files using ABF converter (https://www.reifycs.com/abfconverter/). The .abf files are then transferred to a separate folder which will become the destination folder for the MS-DIAL project. (In MS-DIAL 5, raw files can be directly loaded into the project without conversion).
3. Start MS-DIAL and create a new project by selecting the folder containing all .abf files in *project file path*. The other selections in this window include *Ionisation*: Soft Ionisation; *Separation type*: Chromatography; *MS method*: Conventional LC/MS; *Data type*: Centroid for both MS1 and MS2 (*please ascertain whether the data has been collected in centroid or profile mode by looking into the mass acquisition method used for analysis*); *ion mode*: positive (in our method), *Target omics*: Metabolomics. Click NEXT….
4. In the next window, browse and select all the .abf files under the *Analyse file path* option. It is important here to select the Sample type (Blank/Sample/QC/Standard) and to assign the class ID (e.g. same value for replicates for each sample type). Click NEXT…
5. In the *Analysis Parameter Setting*, there are different tabs with specific features as explained in the table below:

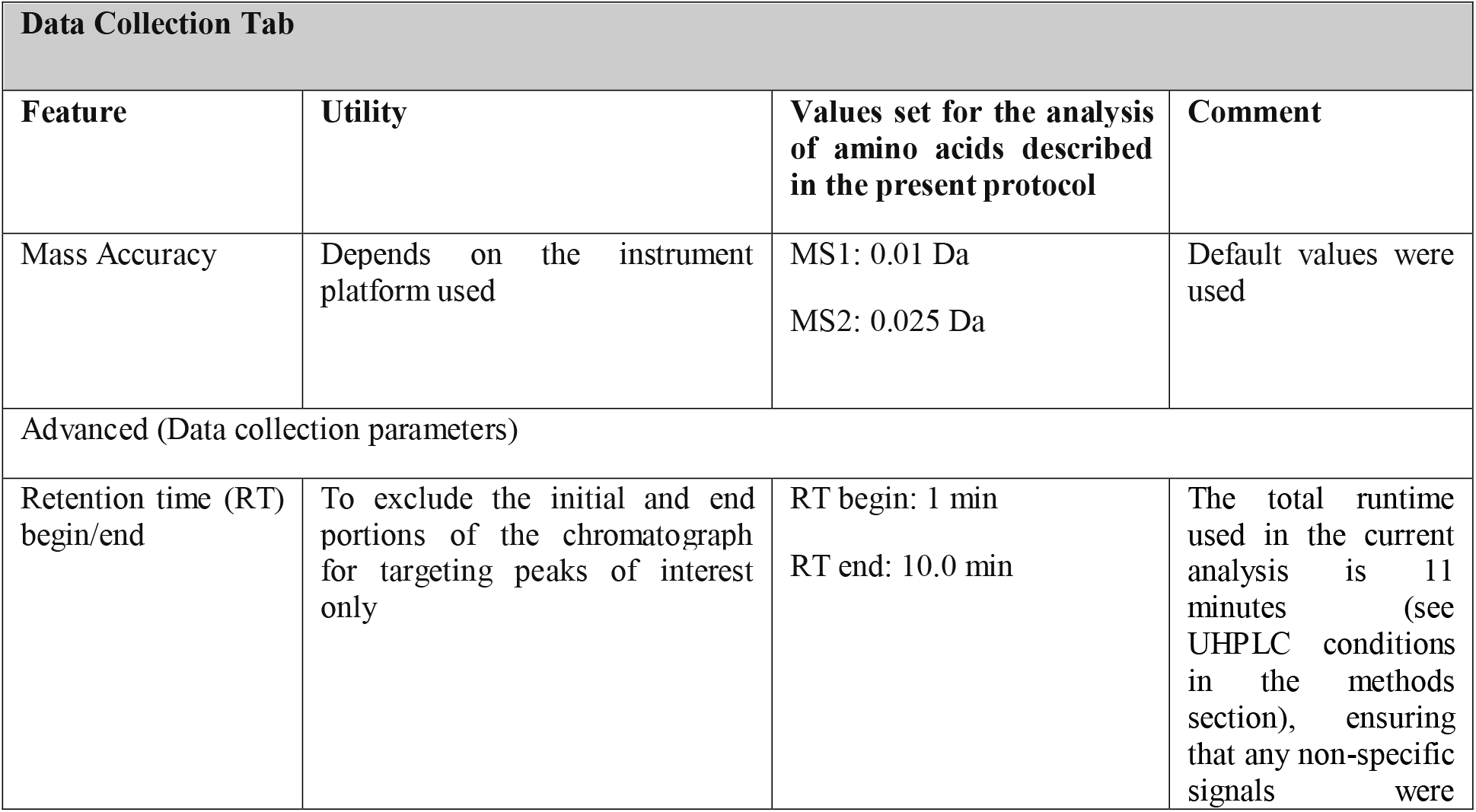
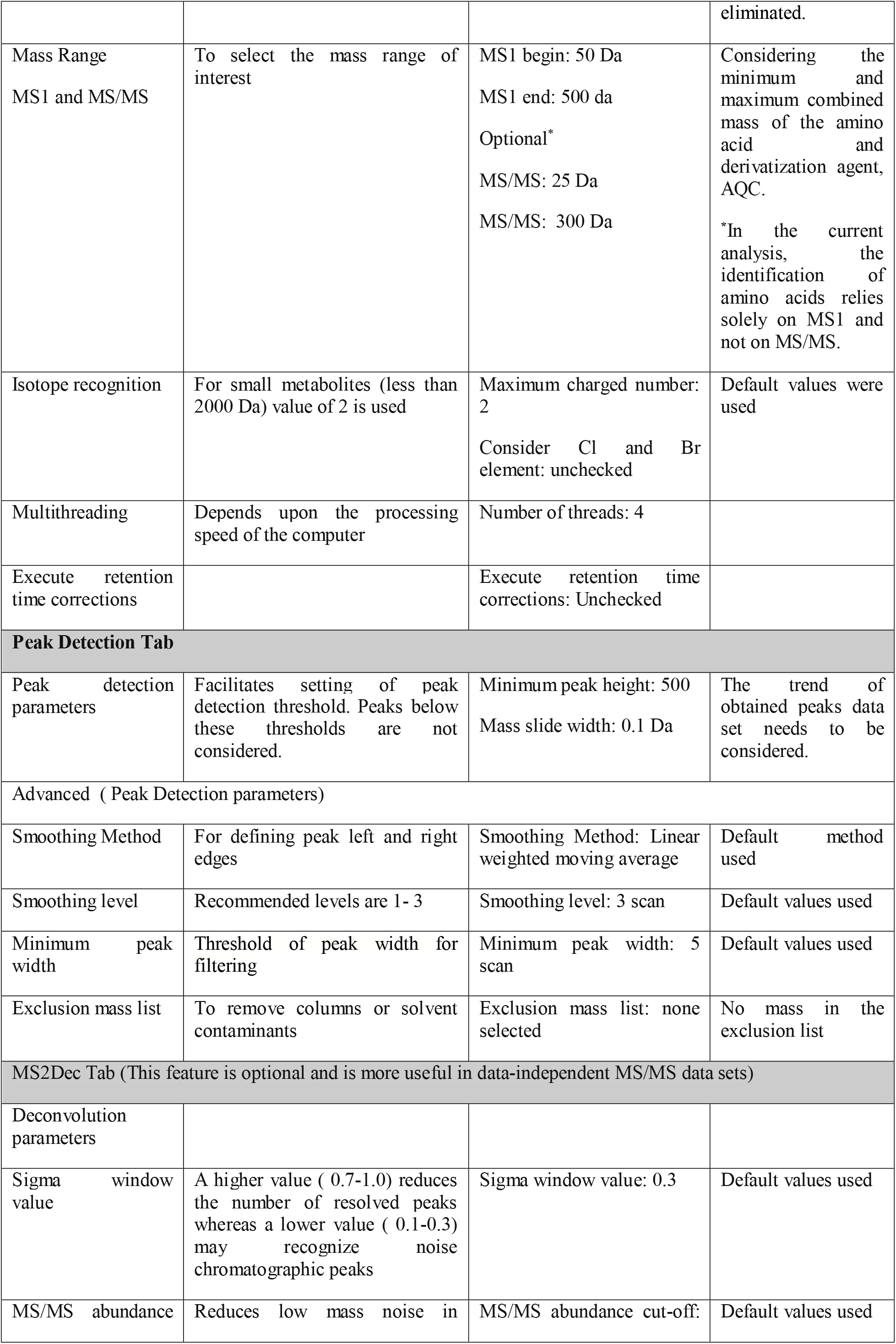
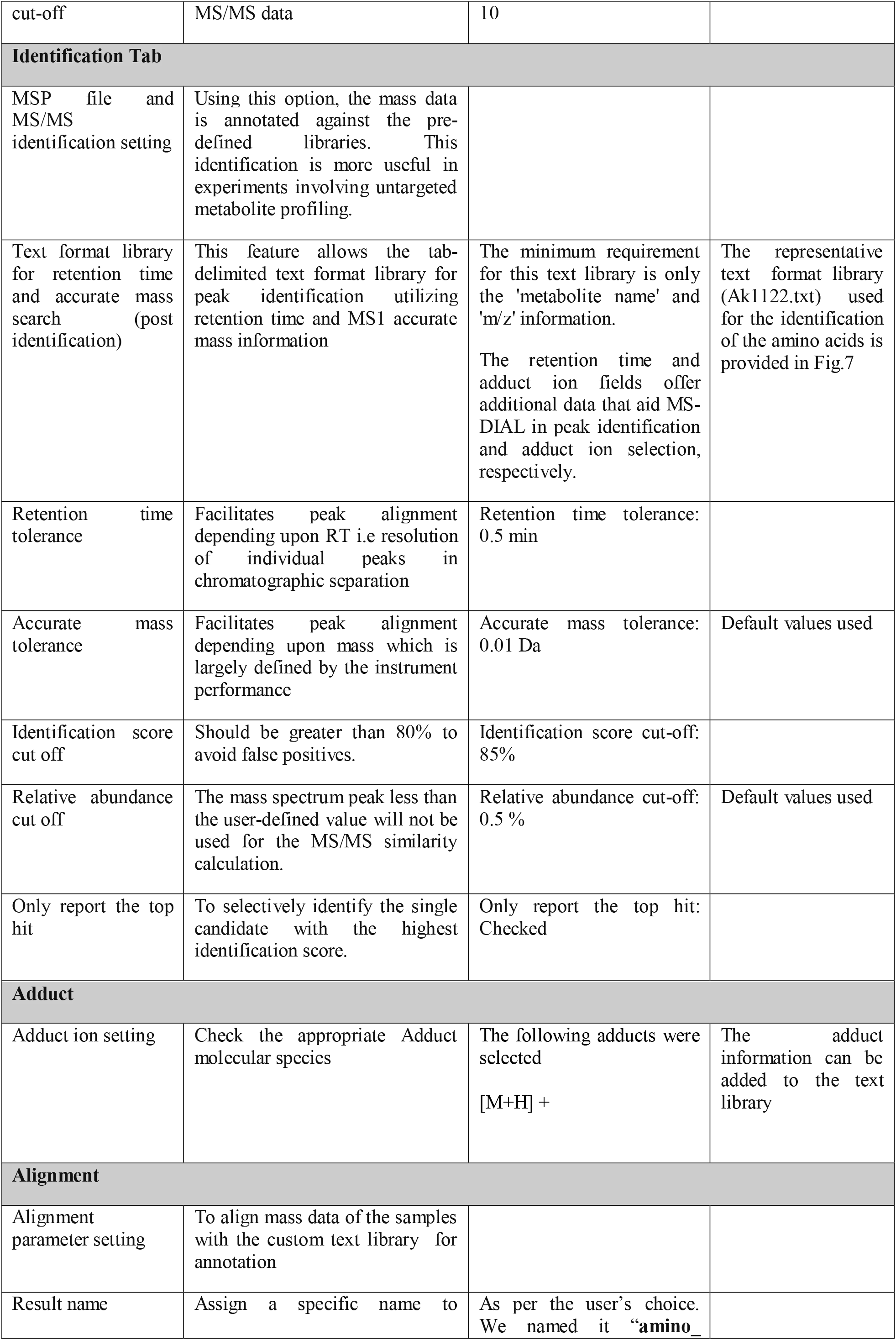
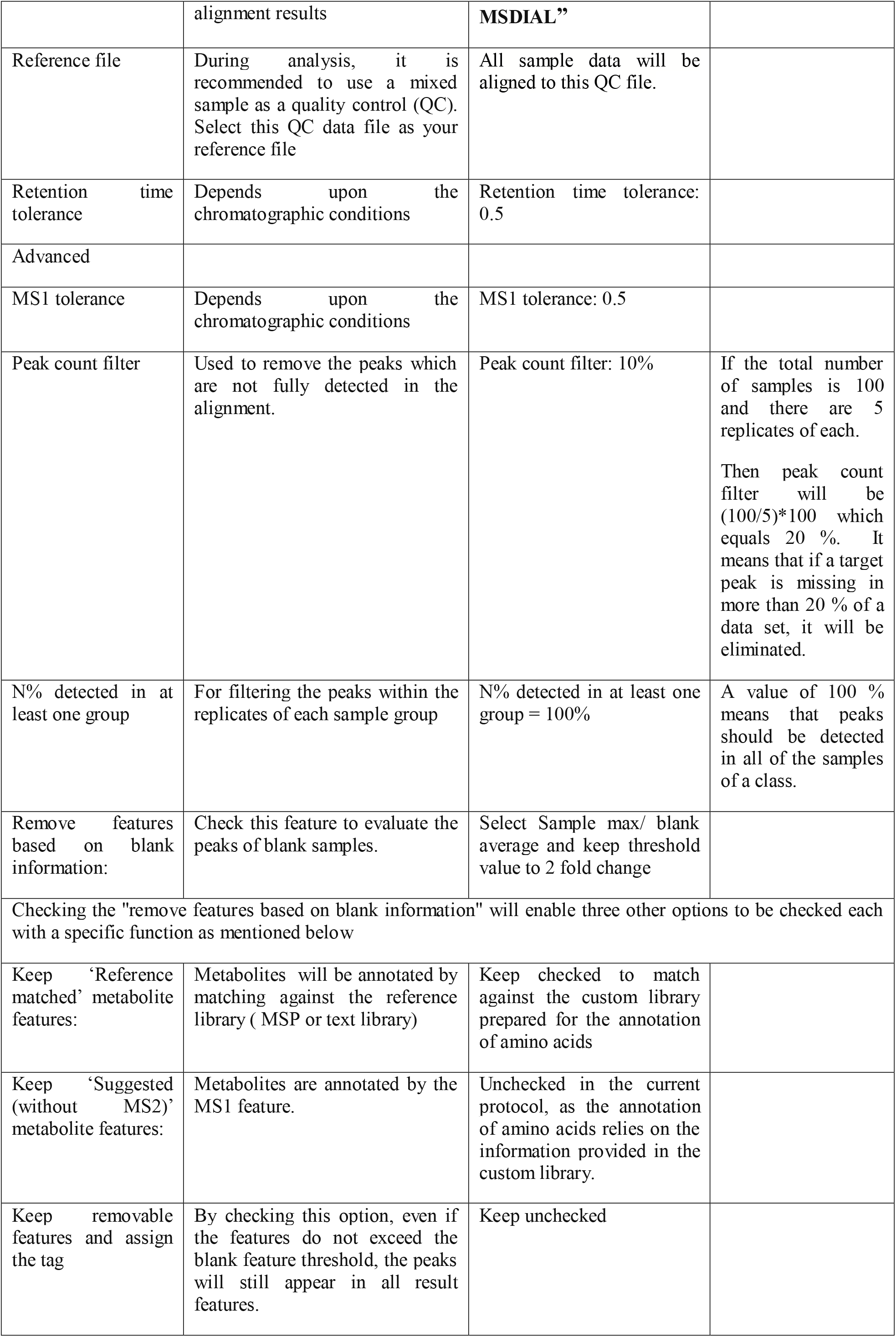
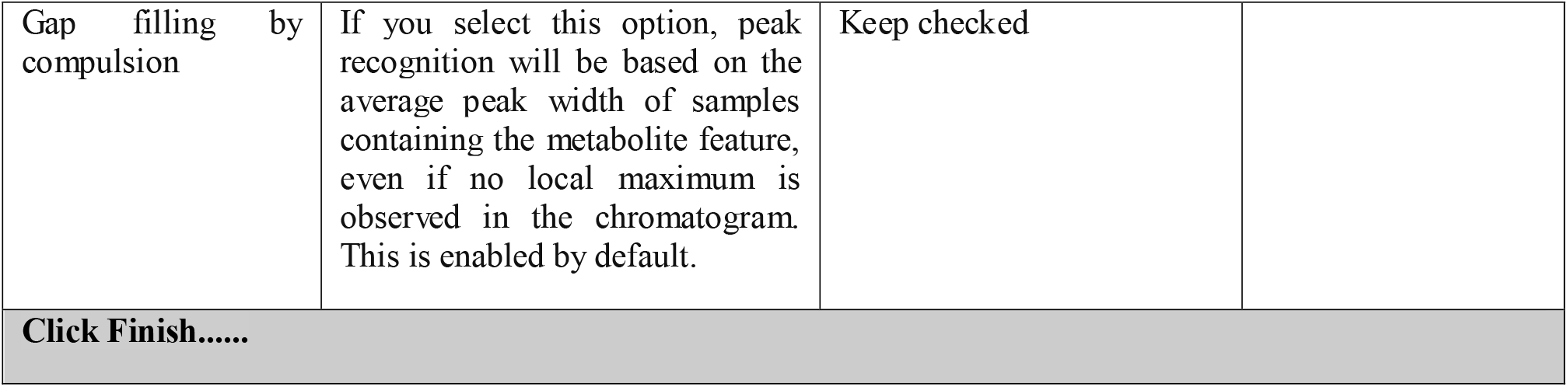

| Data Collection Tab |  |  |  |
| --- | --- | --- | --- |
| Feature | Utility | Values set for the analysis of amino acids described in the present protocol | Comment |
| Mass Accuracy | Depends on the instrument platform used | MS1: 0.01 Da<br>MS2: 0.025 Da | Default values were used |
| Advanced (Data collection parameters) |  |  |  |
| Retention time (RT) begin/end | To exclude the initial and end portions of the chromatograph for targeting peaks of interest only | RT begin: 1 min<br>RT end: 10.0 min | The total runtime used in the current analysis is 11 minutes (see UHPLC conditions in the methods section), ensuring that any non-specific signals were eliminated. |
| Mass Range<br>MS1 and MS/MS | To select the mass range of interest | MS1 begin: 50 Da<br>MS1 end: 500 da<br>Optional*<br>MS/MS: 25 Da<br>MS/MS: 300 Da | Considering the minimum and maximum combined mass of the amino acid and derivatization agent, AQC.<br>*In the current analysis, the identification of amino acids relies solely on MS1 and not on MS/MS. |
| Isotope recognition | For small metabolites (less than 2000 Da) value of 2 is used | Maximum charged number: 2<br>Consider Cl and Br element: unchecked | Default values were used |
| Multithreading | Depends upon the processing speed of the computer | Number of threads: 4 |  |
| Execute retention time corrections |  | Execute retention time corrections: Unchecked |  |
| **Peak Detection Tab** |  |  |  |
| Peak detection parameters | Facilitates setting of peak detection threshold. Peaks below these thresholds are not considered. | Minimum peak height: 500<br>Mass slide width: 0.1 Da | The trend of obtained peaks data set needs to be considered. |
| Advanced ( Peak Detection parameters) |  |  |  |
| Smoothing Method | For defining peak left and right edges | Smoothing Method: Linear weighted moving average | Default method used |
| Smoothing level | Recommended levels are 1- 3 | Smoothing level: 3 scan | Default values used |
| Minimum peak width | Threshold of peak width for filtering | Minimum peak width: 5 scan | Default values used |
| Exclusion mass list | To remove columns or solvent contaminants | Exclusion mass list: none selected | No mass in the exclusion list |
| **MS2Dec Tab (This feature is optional and is more useful in data-independent MS/MS data sets)** |  |  |  |
| Deconvolution parameters |  |  |  |
| Sigma window value | A higher value ( 0.7-1.0) reduces the number of resolved peaks whereas a lower value ( 0.1-0.3) may recognize noise chromatographic peaks | Sigma window value: 0.3 | Default values used |
| MS/MS abundance | Reduces low mass noise in | MS/MS abundance cut-off: | Default values used |
| cut-off | MS/MS data | 10 |  |
| **Identification Tab** |  |  |  |
| MSP file and MS/MS identification setting | Using this option, the mass data is annotated against the pre-defined libraries. This identification is more useful in experiments involving untargeted metabolite profiling. |  |  |
| Text format library for retention time and accurate mass search (post identification) | This feature allows the tab-delimited text format library for peak identification utilizing retention time and MS1 accurate mass information | The minimum requirement for this text library is only the 'metabolite name' and 'm/z' information.<br>The retention time and adduct ion fields offer additional data that aid MS-DIAL in peak identification and adduct ion selection, respectively. | The representative text format library (Ak1122.txt) used for the identification of the amino acids is provided in Fig.7 |
| Retention time tolerance | Facilitates peak alignment depending upon RT i.e resolution of individual peaks in chromatographic separation | Retention time tolerance: 0.5 min |  |
| Accurate mass tolerance | Facilitates peak alignment depending upon mass which is largely defined by the instrument performance | Accurate mass tolerance: 0.01 Da | Default values used |
| Identification score cut off | Should be greater than 80% to avoid false positives. | Identification score cut-off: 85% |  |
| Relative abundance cut off | The mass spectrum peak less than the user-defined value will not be used for the MS/MS similarity calculation. | Relative abundance cut-off: 0.5 % | Default values used |
| Only report the top hit | To selectively identify the single candidate with the highest identification score. | Only report the top hit: Checked |  |
| **Adduct** |  |  |  |
| Adduct ion setting | Check the appropriate Adduct molecular species | The following adducts were selected<br>[M+H] + | The adduct information can be added to the text library |
| **Alignment** |  |  |  |
| Alignment parameter setting | To align mass data of the samples with the custom text library for annotation |  |  |
| Result name | Assign a specific name to | As per the user's choice. We named it "amino_ |  |
|  | alignment results | **MSDIAL”** |  |
| Reference file | During analysis, it is recommended to use a mixed sample as a quality control (QC). Select this QC data file as your reference file | All sample data will be aligned to this QC file. |  |
| Retention time tolerance | Depends upon the chromatographic conditions | Retention time tolerance: 0.5 |  |
| Advanced |  |  |  |
| MS1 tolerance | Depends upon the chromatographic conditions | MS1 tolerance: 0.5 |  |
| Peak count filter | Used to remove the peaks which are not fully detected in the alignment. | Peak count filter: 10% | If the total number of samples is 100 and there are 5 replicates of each.<br>Then peak count filter will be $(100/5)*100$ which equals 20 %. It means that if a target peak is missing in more than 20 % of a data set, it will be eliminated. |
| N% detected in at least one group | For filtering the peaks within the replicates of each sample group | N% detected in at least one group = 100% | A value of 100 % means that peaks should be detected in all of the samples of a class. |
| Remove features based on blank information: | Check this feature to evaluate the peaks of blank samples. | Select Sample max/ blank average and keep threshold value to 2 fold change |  |
| Checking the "remove features based on blank information" will enable three other options to be checked each with a specific function as mentioned below |  |  |  |
| Keep ‘Reference matched’ metabolite features: | Metabolites will be annotated by matching against the reference library (MSP or text library) | Keep checked to match against the custom library prepared for the annotation of amino acids |  |
| Keep ‘Suggested (without MS2)’ metabolite features: | Metabolites are annotated by the MS1 feature. | Unchecked in the current protocol, as the annotation of amino acids relies on the information provided in the custom library. |  |
| Keep removable features and assign the tag | By checking this option, even if the features do not exceed the blank feature threshold, the peaks will still appear in all result features. | Keep unchecked |  |
| Gap filling by compulsion | If you select this option, peak recognition will be based on the average peak width of samples containing the metabolite feature, even if no local maximum is observed in the chromatogram. This is enabled by default. | Keep checked |  |
| Click Finish..... |  |  |  |

The alignment process duration will vary based on the computer’s processing speed. The main outcomes and key post-processing steps are discussed in the results section below. Additionally, a snapshot highlighting the selection of specific features is provided for further clarity

## Results

To obtain the comparative profile of amino acids in different plant samples, click on “**Show ion Table**”. The table contains information on Alignment ID, RT(min), m/z, amino acid name, and the BarChart. In our data, we found considerable modulation in the amino acid levels in control and ToLCPalV-infected *N. benthamiana*. The different time points post-infection are labeled as **1**, 24 h infected; **2**, 24 h control; **3**, 48 h infected; **4**, 48 h control; **5**, 72 h infected; **6**, 72 h control; **7**, 7 d infected; **8**, 7 d control; **9**, 15 d infected; **10**, 15 d control throughout the result section.

Results indicated that 24 hours post-infection, the levels of glycine (Gly), alanine (Ala), threonine (Thr), proline (Pro), glutamic acid (Glu), histidine (His), phenylalanine (Phe), and tyrosine (Tyr) increased in infected plants compared to the control plants. Although amino acid levels increased in both infected and control plants after 72 hours of infection, infected plants exhibited a higher accumulation of most amino acids compared to control plants. After 7 days post-infection, proline (Pro), histidine (His), and tyrosine (Tyr) levels remained higher in infected plants compared to control plants. Notably, after 15 days post-infection, the levels of most amino acids decreased in infected plants as compared to the control plants (Fig. 1)

**Fig.1:**
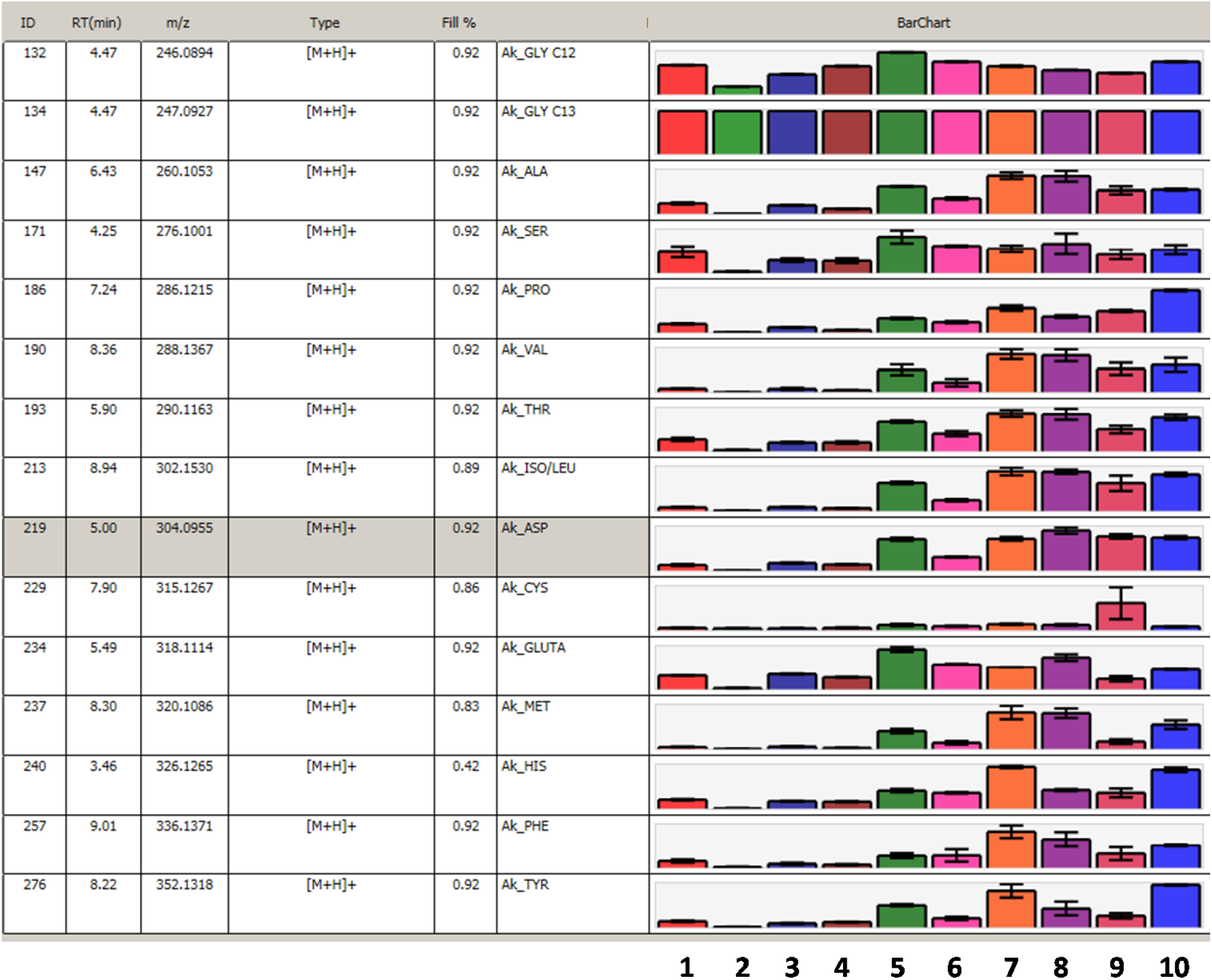
The result obtained from MS-DIAL showing significant modulation in the levels of amino acids in control and ToLCPalV-infected *N. bethamiana* plants at different time points post-infection. 1, 24 h infected; 2, 24 h control; 3, 48 h infected; 4, 48 h control; 5, 72 h infected; 6, 72 h control; 7, 7 d infected; 8, 7 d control; 9, 15 d infected; 10, 15 d control.

The MS-DIAL output window includes several sections, such as file navigator, alignment navigator, peak spot navigator, MS1 spectrum, EIC of focused, aligned spot, bar chart of aligned spot, peak/aligned spot viewer, show ion table, compound detail, and experiment vs reference spectrum viewer as shown in Fig. 2

**Fig. 2:**
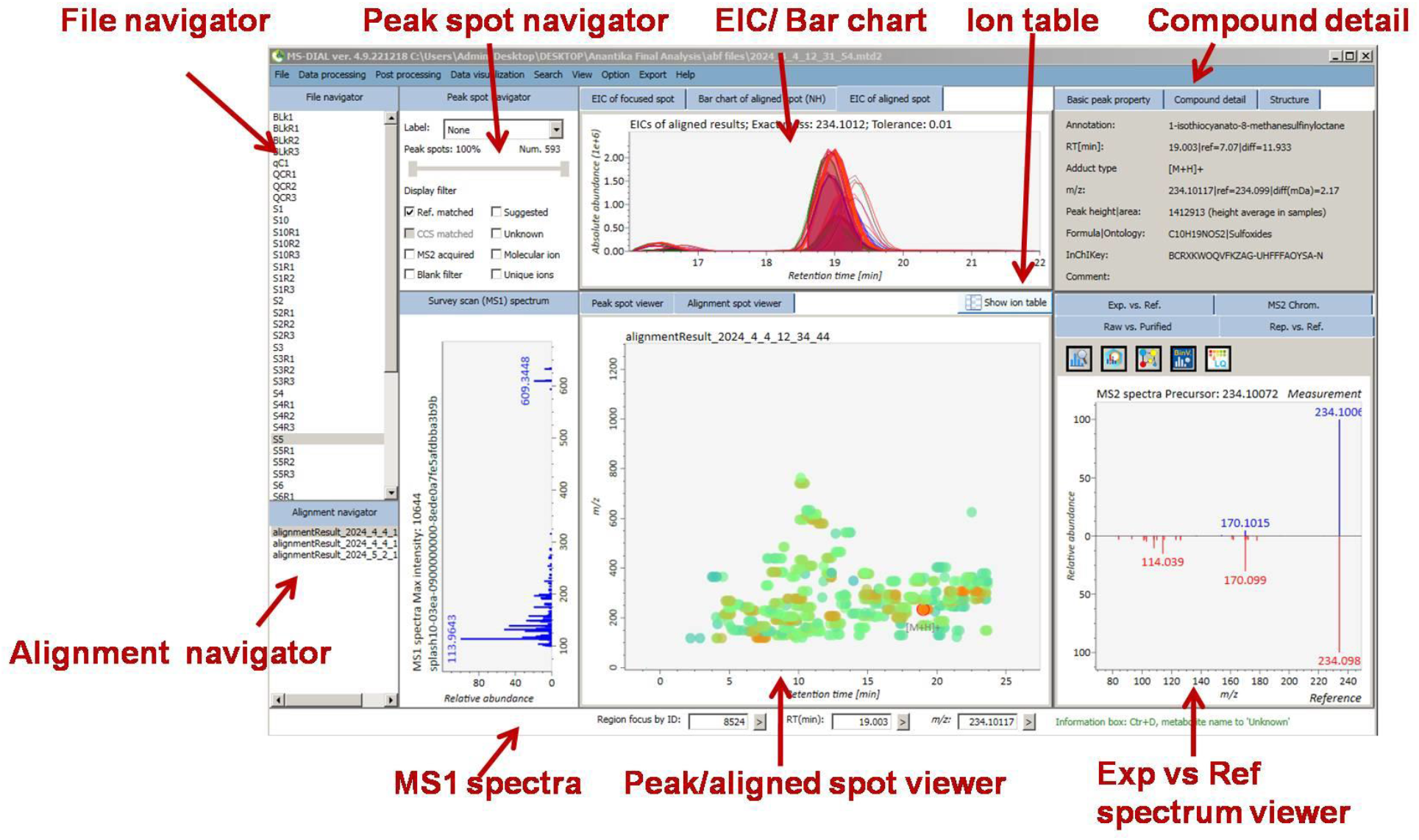
Picture showing different navigator windows present in MS-DIAL.

The alignment results were obtained by clicking on the alignment name (amino _MSDIAL) in the alignment navigator. To display only the aligned spots that match those in the custom text library, it is important to select “Ref matched” in the peak spot viewer. The results showed that a total of 15 amino acids were identified across the 10 different samples (control and virus-infected tobacco plants) used in this study. At this stage, various features, including the EIC of aligned spots, MS1 spectrum, and Exp vs Ref spectrum, can be explored to ensure the accuracy of the alignment. Notably, selecting the “bar chart of aligned spot” will display the relative abundance of amino acids present in the dataset under study. By clicking on each amino acid aligned spot, the respective bar chart will appear in the section “bar chart of aligned spot” (Fig. 3).

**Fig. 3:**
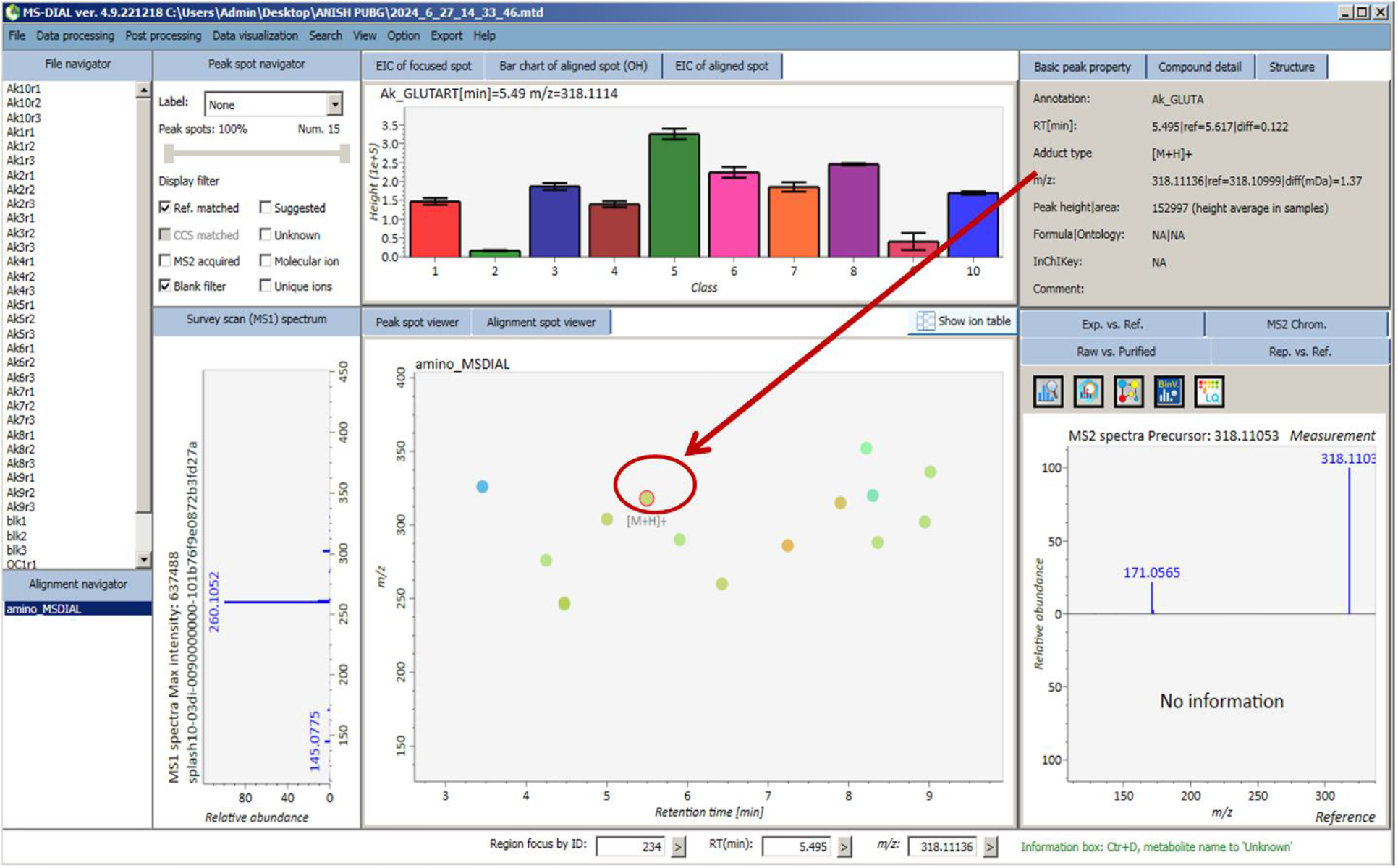
Alignment results showing the selection of spots corresponding to glutamate (m/z of 318.11 and RT of 5.617) and its relative levels in control and ToLCPalV -infected plants at different time points of infection.

It is important to note that variations in sample preparation during extraction and/or chemical derivatization can affect the relative levels of amino acids in different samples. Therefore, normalizing the dataset is crucial for obtaining an accurate relative profile of amino acids. MS-DIAL provides various methods for data normalization, including internal standard, LOWESS, internal standard + LOWESS, SPLASH lipidomix (for lipidomics), Total Ion Chromatograms (TIC), and TIC of identified metabolites (mTIC). In the present method, we will focus exclusively on internal standard-based data normalization, Glycine-1-13C (1 µg/ml) was added during the extraction of amino acids.

- Select the target ID of glycine-1-13C (internal standard) in the “normalization property setting” for alignment results.
- In our results, the alignment ID of glycine-1-13C with a mass of 247.0921 was 134. The same value needs to be filled in the Target ID column for each row as shown in the Fig. 4.
- For normalisation, click ‘normalisation” from the “Data visualisation” tab. Select the option of internal standard and click Done.
- Click Finish. A clear difference in the “bar chart of aligned spot” can be observed before and after normalization (Fig.4).

**Fig. 4:**
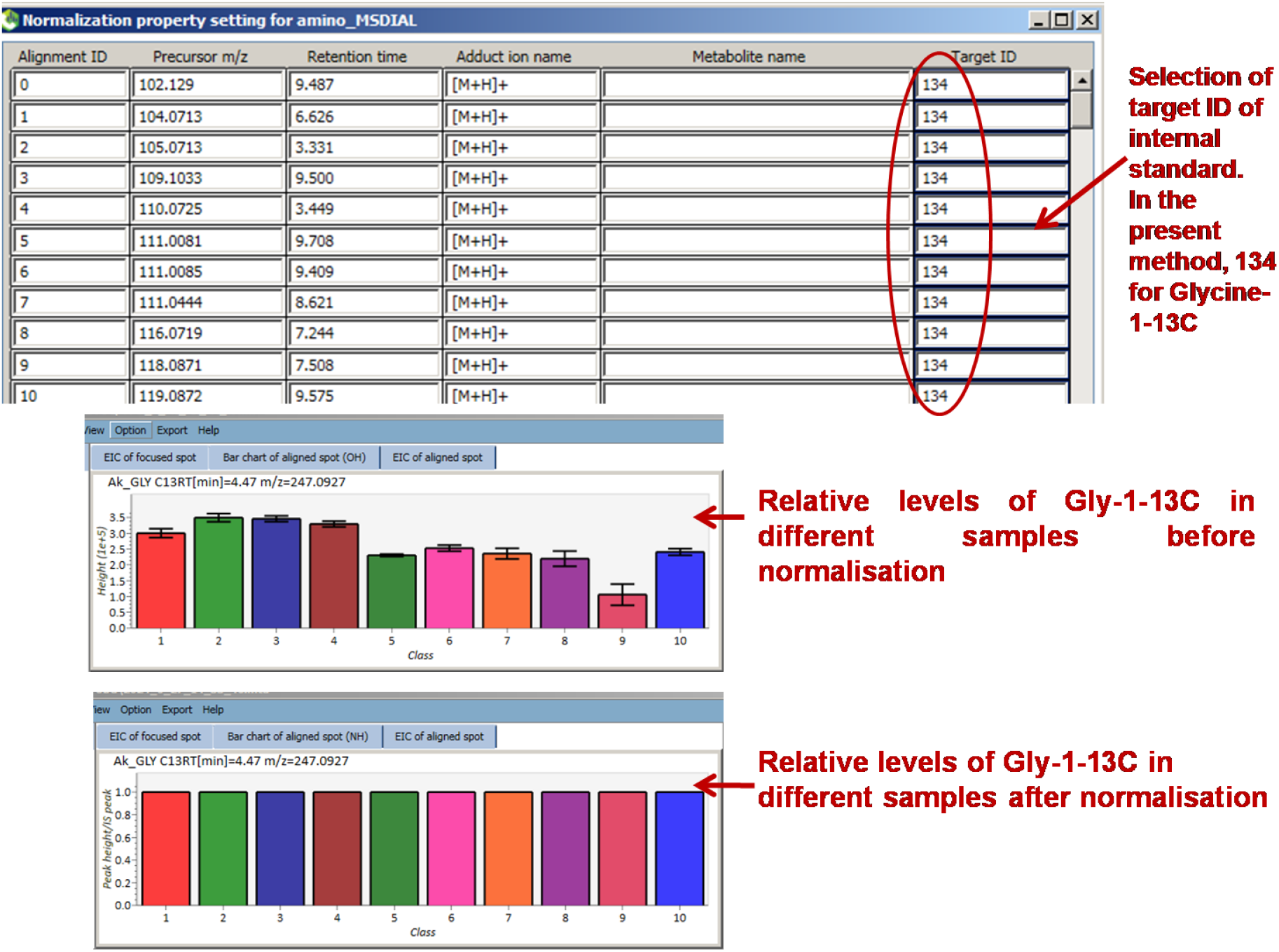
Alignment ID (134) of Gly-1-13C selected for each spot in the Target ID column. The lower panel shows the relative levels of Gly-1-13C in different samples before and after normalization.

Further, MS-DIAL provides an option to curate mass peaks, which are then saved and displayed in the “bar chart of aligned spots.” This feature is quite useful to determine whether the variation within replicates of the same sample is occurring naturally or is due to incorrect integration of mass peak. At first, the samples showing higher variation need to be identified. For this, click on the “ Show ion Table “. Notably, in our data, none of the amino acids showed large variation due to incorrect integration of mass peak. Therefore, we selected the mass peak of “cys” from the previous data set for demonstration as detailed below:

Select “ EIC of aligned spot “ adjacent to the “ Bar chart of aligned spot “. Right-click on the EIC region and select “Table viewer for curating each chromatograph “. Select the target chromatograph and manually integrate the peak by dragging the red dot (Fig. 5). Select update and observe the change in color for peak intensity and peak area in the Table viewer.

**Fig. 5:**
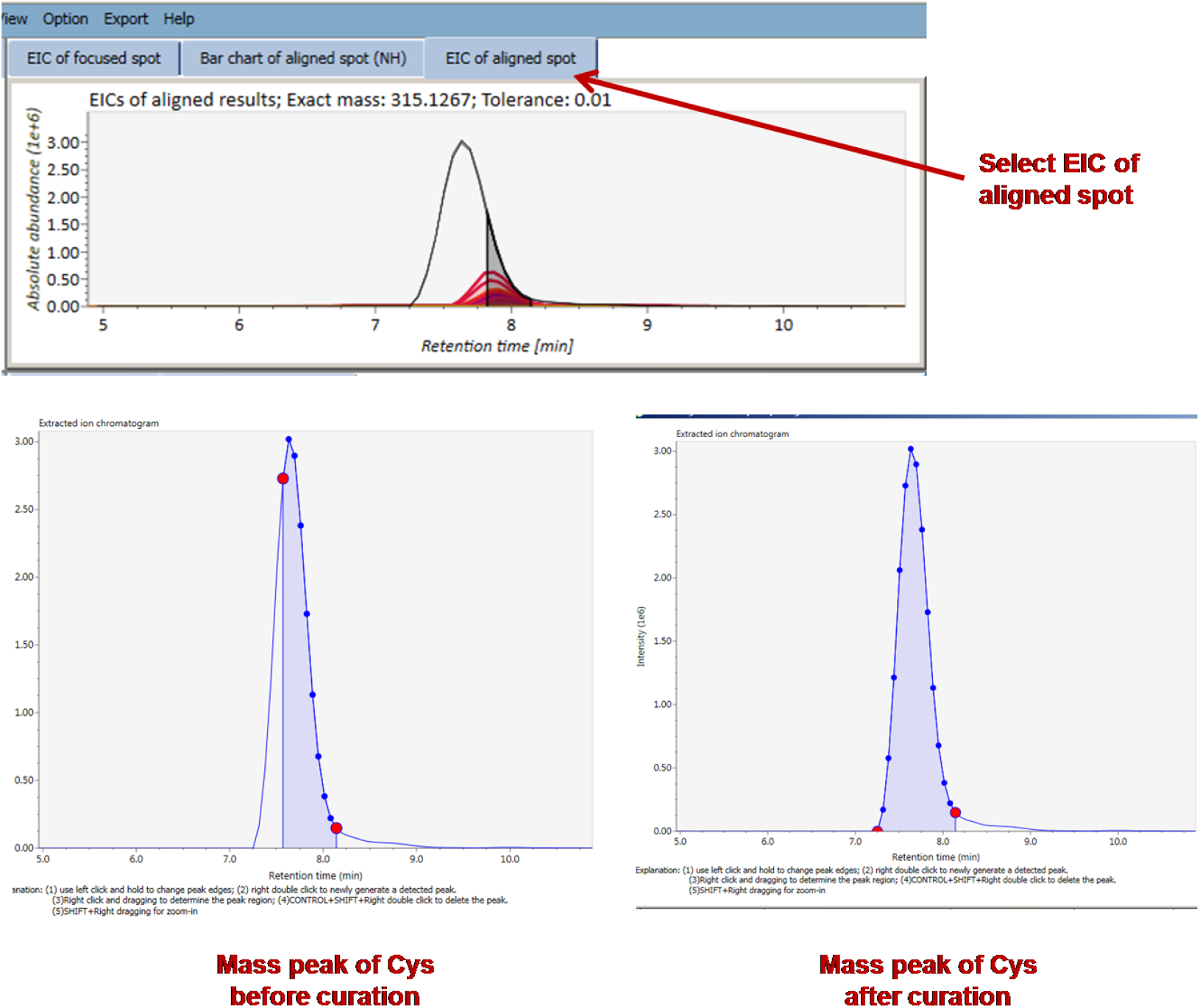
Curation of an incorrectly integrated mass peak.

Another important feature is Principal Component Analysis (PCA) which can be selected from the “Data Visualization” tab. The PCA defines the statistical variation among different samples and is hence important to infer variation amongst large data sets.

Click PCA from “Data visualization” tab.

1. A PCA setting window will pop up. In this window fill maximal principal component: 5; Scale method: auto scale; Transform method: none; Metabolite selection: Ref matched. These values depend upon the complexity of the data sets and are selected accordingly.
2. Click Done to obtain the PCA Plot. The results are shown in Fig. 6

**Fig. 6:**
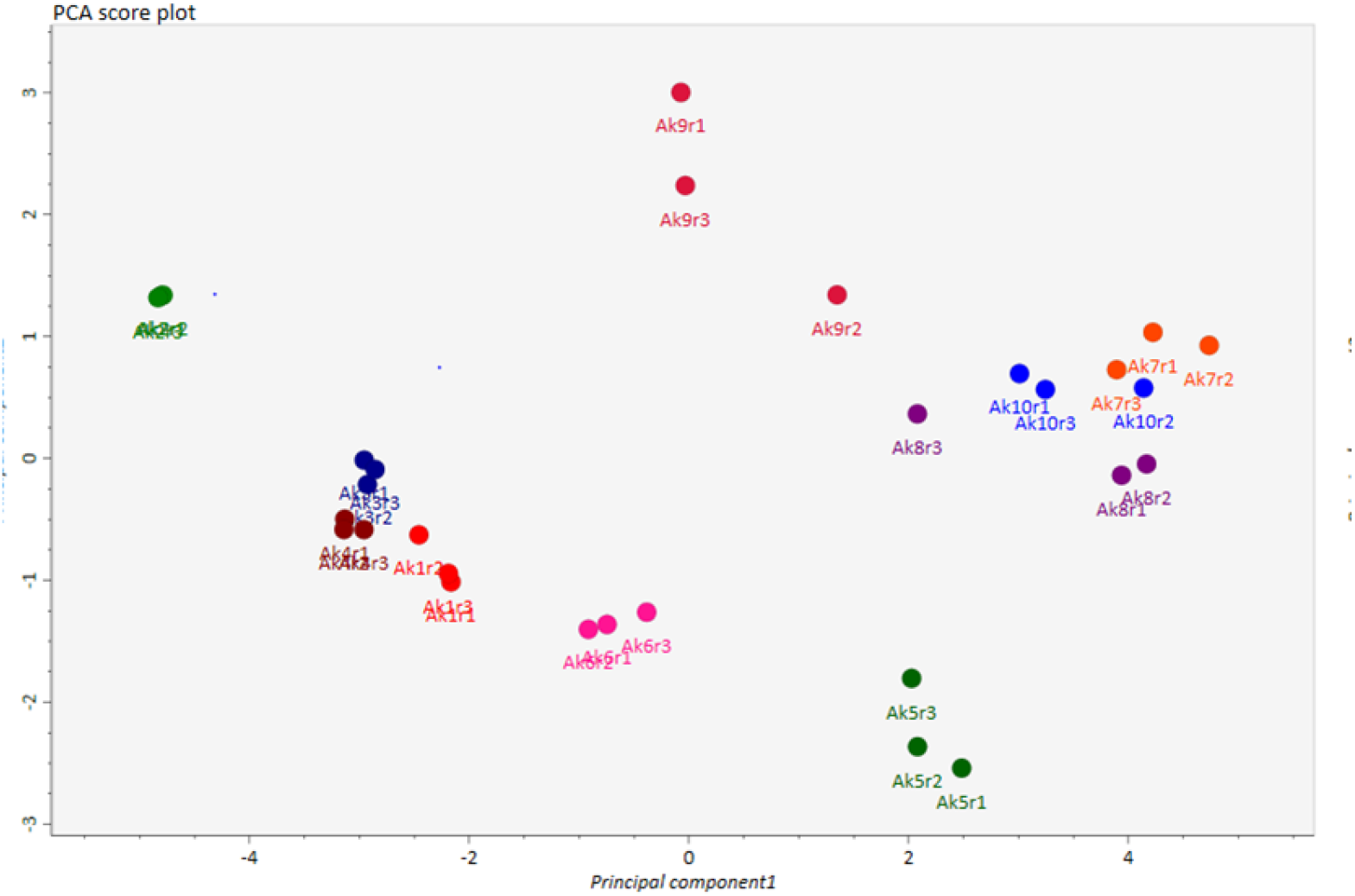
Principal component analysis (PCA) score plot showing significant difference exists in the levels of amino acids in control and ToLCPalV -infected plants at different time points of infection. 1, 24 h infected; 2, 24 h control; 3, 48 h infected; 4, 48 h control; 5, 72 h infected; 6, 72 h control; 7, 7 d infected; 8, 7 d control; 9, 15 d infected; 10, 15 d control.

**Fig. 7:**
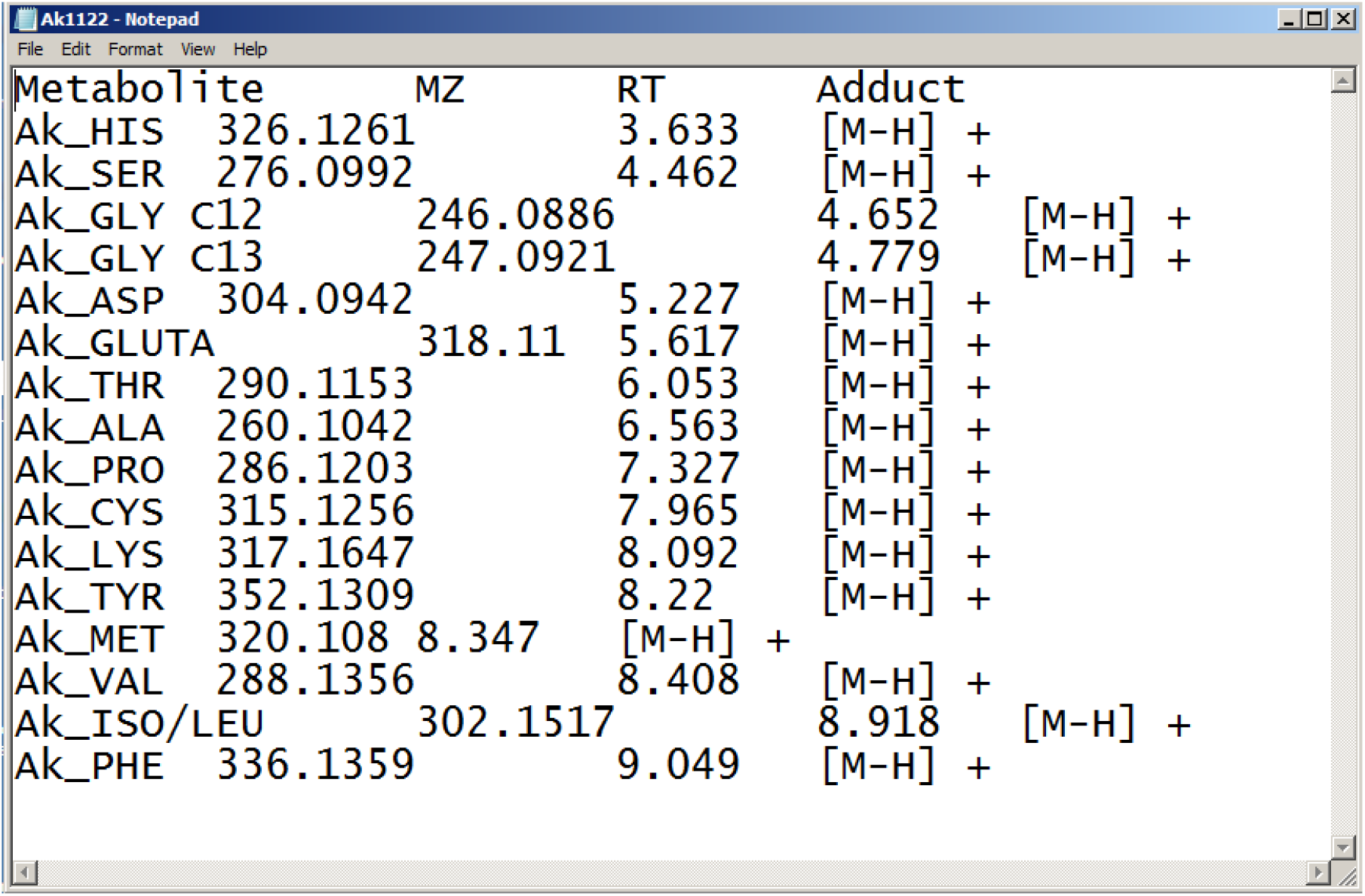
A snapshot of the library in text format containing information on m/z and retention time (RT) of individual amino acids. This custom library is used for post-run identification of amino acids as detailed in the method section.

## Conclusion

Estimating relative levels of amino acids is crucial for understanding various biological processes in plants. The present method provides a detailed protocol using MS-DIAL for obtaining a qualitative amino acid profile from LC-MS data. The main objective of the present method is to simplify data processing and enhance confidence in assessing relative changes in the levels of amino acids. Aligning the mass peak data with a custom library carrying information on both m/z and RT of different amino acids significantly enhances the accuracy of peak identification and annotation. Importantly, the workflow detailed in this protocol can be easily adapted for the analysis of other metabolites for which reference standards are available.

## Other Useful Resources

https://www.researchgate.net/publication/359128438_MSDIAL_for_dummies_An_honest_tutorial_for_MS-DIAL_use_in_metabolomics_data_analysis

https://www.youtube.com/watch?v=yPawtF1qJ5E

https://www.youtube.com/@msdialproject

